# Preclinical assessment of CAR-NK cell-mediated killing efficacy and pharmacokinetics in a rapid zebrafish xenograft model of metastatic breast cancer

**DOI:** 10.1101/2023.07.11.548344

**Authors:** Nivedha Murali Shankar, Paola Ortiz Montero, Anastasia Kurzyukova, Wiebke Rackwitz, Stephan R. Künzel, Winfried S. Wels, Torsten Tonn, Franziska Knopf, Jiri Eitler

## Abstract

Natural killer (NK) cells are attractive effectors for adoptive immunotherapy of cancer. Results from first-in-human studies using chimeric antigen receptor (CAR)-engineered primary NK cells and NK-92 cells are encouraging in terms of efficacy and safety. In order to further improve treatment strategies and to test the efficacy of CAR-NK cells in a personalized manner, preclinical screening assays using patient-derived tumor samples are needed. Zebrafish (*Danio rerio*) embryos and larvae represent an attractive xenograft model to study growth and dissemination of patient-derived tumor cells because of their superb live cell imaging properties. Injection into the organism’s circulation allows investigation of metastasis, cancer cell-to-immune cell-interactions and studies of the tumor cell response to anti-cancer drugs.

Here, we established a zebrafish larval xenograft model to test the efficacy of CAR-NK cells against metastatic breast cancer *in vivo* by injecting metastatic breast cancer cells followed by CAR-NK cell injection into the Duct of Cuvier (DoC). We validated the functionality of the system with two different CAR-NK cell lines specific for PD-L1 and ErbB2 (PD-L1.CAR NK-92 and ErbB2.CAR NK- 92 cells) against the PD-L1-expressing MDA-MB-231 and ErbB2-expressing MDA-MB-453 breast cancer cell lines.

Injected cancer cells were viable and populated peripheral regions of the larvae, including the caudal hematopoietic tissue (CHT), simulating homing of cancer cells to blood forming sites. CAR-NK cells injected 2.5 hours later migrated to the CHT and rapidly eliminated individual cancer cells throughout the organism. Confocal live-cell imaging demonstrated intravascular migration and real-time interaction of CAR-NK cells with MDA-MB-231 cells, explaining the rapid and effective *in vivo* cytotoxicity.

Thus, our data suggest that zebrafish larvae can be used for rapid and cost-effective *in vivo* assessment of CAR-NK cell potency and to predict patient response to therapy.

## INTRODUCTION

Retargeting of T cells with CAR against a tumor marker has shown remarkable clinical efficacy in relapsed or refractory B-cell malignancies and multiple myeloma (Finck et al., 2022). This success has led to the approval of several CAR T cell products targeting hematologic malignancies by the U.S. Food and Drug Administration (FDA) and the European Medicines Agency (EMA). Despite significant clinical efficacy in some cancers, CAR T cell therapy is often associated with several side effects, such as a cytokine storm or neurological toxicity. In addition, CAR T cell products must be produced autologously due to T cell product incompatibility between patients, resulting in high therapy costs. As an alternative, NK cells have recently been recognized as a promising cytotoxic cell type for CAR-based cancer therapy, overcoming many of the obstacles of T cell-based systems. Primary NK cells and the NK cell line NK-92 have been shown to be safe with no adverse effects in clinical trials, and early clinical data in hematologic malignancies are encouraging in terms of efficacy (Burger et al., 2023; Liu et al., 2020; Tonn et al., 2013). NK cells do not express the T cell receptor and therefore an NK cell product can be infused into multiple recipients in an allogeneic scenario without the risk of graft versus host disease (GvHD). In contrast, NK cells express an inherited set of activating or inhibitory receptors that allow them to recognize tumor or infected cells, making them unique cytotoxic cells. Although NK cells can recognize and kill many cancer cells, some cancers can evade NK cell-mediated killing. To overcome this resistance, NK cells can also be genetically engineered with CAR, similar to T cells (Romanski et al., 2016; Schonfeld et al., 2015; Uherek et al., 2002). Indeed, several sources of NK cells are in preclinical or clinical development, including primary blood-derived NK cells, NK cells differentiated from hematopoietic or induced pluripotent stem cells (Liu et al., 2020; Miller et al., 2005; Zhu et al., 2018). In addition, stable NK cell lines such as NK-92 have been established for clinical use (Nowakowska et al., 2018; Tonn et al., 2013).

Despite the success of CAR-based therapies in hematological malignancies, solid tumors such as breast cancer remain a challenge. CAR T cells have not shown clinical efficacy and CAR-NK cells are currently in clinical investigation. The difficulties in targeting solid tumors are determined by inter-tumor heterogeneity, heterogeneity in antigen expression, resistance mechanisms to CAR T or NK cells and often generate metastatic diaspores that are difficult to reach by immune cells. Therefore, a personalized approach is required to select the most appropriate CAR system for each patient. Novel CAR-based strategies are rapidly developing and good preclinical models are needed to apply them in a personalized manner. Currently, novel CAR-NK cell therapies are tested in immunodeficient mouse models. However, these models are very expensive, laborious, inflexible and slow to yield results. Furthermore, the lack of transparency of the host tissues hampers *in vivo* observations of homing processes and the antitumor response. Therefore, in order to bridge the gap from *in vitro* studies to clinical trials, medium to high-throughput preclinical models with optimal live-cell imaging properties and fast experimental readout are needed to test the efficacy of CAR-NK cells against patient cancer cells.

Zebrafish (*Danio rerio*) embryos and larvae are prime candidates for the development of such a xenograft screening system. Their small size and transparency allow for medium-throughput injections with the prospect of automated high-throughput upscaling in the near future. Likewise, imaging of cancer and immune cells in transplanted embryos and larvae can be performed at high throughput at low cost. Notably, the lack of a fully functional adaptive immune system during early larval stages prevents rejection of transplanted cells in zebrafish (Lam et al., 2004). Zebrafish xenografts have been developed for the investigation of human tumor cell proliferation, apoptosis, invasion, extravasation and small molecule anti-cancer drug screening (Dietrich et al., 2021; Miao et al., 2021). Recent work has shown that the zebrafish model can be used to assess the *in vivo* efficacy of CAR T cells (He et al., 2020; Pascoal et al., 2020; Zhou et al., 2022). This provides proof of principle that embryonic zebrafish xenografts can be used for preclinical testing of CAR T cell efficacy; however, whether zebrafish are suitable to evaluate the antitumor action of CAR-NK cells is currently unclear.

Here, we established zebrafish embryos and larvae as a preclinical xenograft model to assess the efficacy of CAR-NK cells against metastatic cancer cells of solid tumor origin. We validated the CAR-NK cell system with two different specificities and two different breast cancer models. Furthermore, we investigated the live interaction of CAR-NK and cancer cells and the migration of CAR-NK cells through the microvasculature to the site of metastatic breast cancer cells. This demonstrates the functionality and usefulness of CAR-NK cells in cancer therapy and reaffirms the power of zebrafish xenografts in preclinical cancer research.

## MATERIALS AND METHODS

### Cells and cell culture

The established human NK cell line NK-92 was provided by H.G. Klingemann (Chicago, IL, USA) (Maki et al., 2001). ErbB2.CAR NK-92 (NK-92/5.28.z) cells were generated previously (Schonfeld et al., 2015). NK cell lines were cultured in X-vivo 10 medium (Lonza) containing 5% heat-inactivated human AB plasma (German Red Cross Blood Donation Service North-East, Dresden, Germany), 500 IU/mL IL-2 (Proleukin; Novartis Pharma), 100 IU/mL penicillin, and 100 µg/mL streptomycin (Merck/Biochrom). MDA-MB-453, MDA-MB-231 cells were purchased from the American Type Culture Collection (ATCC, Manassas, VA, USA) and cultured in DMEM medium (Merck/Biochrom) supplemented with 10% HI-FBS (Merck/Biochrom), 2 mM L-glutamine (Merck/Biochrom), 100 IU/mL penicillin, and 100 μg/mL streptomycin. All cells were cultured at 37°C in a humidified atmosphere with 5% CO_2_ and routinely checked for *Mycoplasma* contamination.

### Generation of transgenic cells

PD-L1.CAR NK-92 were generated by lentiviral transduction with a second-generation CAR consisting of PD-L1-specific scFv, CD8 hinge, CD28 transmembrane/costimulatory, and CD3ζ intracellular signaling domains cloned into the pSIEW backbone as described previously (Schonfeld et al., 2015). PD-L1.CAR NK-92 cells were further immuno-magnetically enriched according to manufacturer’s protocol (Miltenyi Biotec) using human recombinant PD-L1-Fc combined with biotinylated anti-human Fc. Enriched cells resulted in >95% purity of PD-L1.CAR-expressing NK-92 cells at the time of analysis. MDA-MB-231 and MDA-MB-453 cells were transduced with pSIEW-GFP/Puro vector at MOI<1 by 30 min spinoculation at 1000xg in the presence of 8µg/ml Polybrene (Sigma-Aldrich) to produce MDA-MB-231 GFP and MDA-MB-453 GFP. Cells were purified by puromycin selection (Sigma-Aldrich).

### Lentivirus production

Lentivirus particles were produced in the packaging cell line HEK293T using packaging vectors psPAX2 and pMD2.G (both from Addgene). Plasmids were transfected with polyethyleneimine (PEI) and supernatants were harvested after 48 hours. Virus was concentrated with PEG-it solution (System Biosciences) according to the manufacturer’s instructions.

### Flow cytometry

Cells were stained with antibodies specific for PD-L1 (Miltenyi Biotec) and ErbB2 (R&D Systems) for 30 min on ice. For CAR expression, CAR-NK cells were loaded with human recombinant PD-L1- Fc (Biolegend) or ErBb2-Fc fusion proteins (R&D Systems) for 30 min on ice, washed, and stained with anti-human Fc secondary antibody (Jackson Immunoresearch) for 30 min on ice. Live cells were discriminated using 7-AAD (BD Biosciences) or DAPI (Sigma Aldrich). Samples were acquired using a BD FACSCanto II flow cytometer and data were analyzed using FlowJo software version 9 (BD Biosciences).

### Europium-TDA (EuTDA) cytotoxicity assay

The specific cytotoxicity of the NK-92 cell lines against target cells was determined using the Europium (EuTDA) cytotoxicity assay (DELFIA, PerkinElmer #C135-100) according to the manufacturer’s protocol and as previously described (Eitler et al., 2021). Briefly, target cells were loaded with an acetoxymethyl ester of the fluorescence-enhancing ligand (BATDA; Perkin Elmer #C136-100) and then co-cultured at 10,000 cells/well in triplicate with effector cells at the indicated E:T ratios. After 2 hours of co-culture, supernatants were collected for measurement of the fluorescent signal reflecting target cell lysis using a VICTOR X4 fluorometer (PerkinElmer). Specific lysis was calculated according to the standard formula in the manufacturer’s instructions.

### Zebrafish husbandry

Zebrafish were maintained in the fish facility of the Center for Regenerative Therapies, Dresden (CRTD) in accordance with the guidelines of the Landesdirektion Dresden (DD25-5131/450/4 and 25- 5131/564/2). The experiments were performed with zebrafish larvae up to a maximum age of 5 dpf (days post fertilization). Adult fish were mated in mating tanks and the eggs were collected 2-3 hours after mating. Fertilized eggs were raised in E3 medium (5 mM NaCl, 0.17 mM KCL, 0.33 mM CaCl2, 0.33 mM MgSO4) at 28°C.

### Cell preparation and labelling

At 2 dpf, MDA-MB-231 GFP or MDA-MB-453 GFP cells were detached using Accutase (Biolegend) and concentrated to 1×10^6^ cells/ml in DMEM. NK-92 or CAR NK-92 cells were labeled with PKH26 (Sigma Aldrich) according to the manufacturer’s instructions. Briefly, 2×10^6^ NK cells were washed with serum-free X-vivo medium followed by resuspension in 200 µl diluent C (DC), mixed with PKH26 staining solution (1µl PKH26 in 250 µl DC) and incubated for 5 min at RT. The reaction was stopped by adding FBS for 1 min. The labeled cells were washed twice with pure X-vivo medium and finally concentrated to 1×10^6^ cells/ml in X-vivo medium. The cells were validated for staining by flow cytometry. Shortly before injection, cancer and NK cells were washed with 1X PBS, followed by Tx buffer (1% Pen/Strep, 1.5 mM EDTA, 1X PBS) and finally resuspended in 10 µl of Tx buffer (1×10^6^ cells/10 µl, 100 cells/nl) and kept on ice.

### Microinjection

Wild type (WT) AB or transgenic Tg(*flk1*:GFP) zebrafish embryos were dechorionated at 2 dpf using forceps (Dumostar Biology, Dumont 55), anesthetized in 0.02% tricaine methanesulfonate (MESAB; Sigma Aldrich #A5040) in E3, aligned on 1.5% agarose plates with grooves, and placed under a dissecting microscope (Olympus SZX10) equipped with a microinjector (Marzhauser, MM3301R) and a vacuum pump (WPI, Pneumatic PicoPump PV 820). Filamented glass needles (TW100F-3, World Precision Instruments) pulled with a P-97 Flaming/Brown micropipette puller (Sutter Instrument) were used to inject 1 nl (in average 100 cells) of cancer cells into the Duct of Cuvier (DoC), followed by fresh addition of E3 to the injected larvae and incubation at 33 □C to allow recovery. After 2.5 hours, larvae were sorted by fluorescence microscopy to discard improperly injected larvae. The second microinjection was performed to inject 1 nl of labeled NK cells followed by E3 addition and incubation at 33 □C.

### Zebrafish imaging

For imaging, larvae were anesthetized in 0.02% MESAB in E3, embedded in 2.5% methylcellulose in a glass cavity slide and imaged using the SteREO Discovery V12 microscope equipped with AxioCam MRm (Zeiss) and AxioVision 4.7 software (Zeiss). Fluorescence imaging was performed on both sides of the larvae in brightfield, GFP and RFP channels, wherein unsaturated images were obtained.

### Confocal Microscopy

Zebrafish larvae [WT AB and Tg(*flk1*:GFP)] were injected with MDA-MB-231 GFP followed by PKH26-labeled PD-L1.CAR NK-92 cells. For confocal imaging, larvae were anesthetized with 0.02% MESAB in E3 and embedded in 1% low melt agarose (LMA) filled with 0.01% MESAB in E3. High-resolution live imaging of larvae was performed using a Dragonfly spinning disk confocal microscope (Andor) equipped with a sCMOS camera, and Fusion software (Andor, version 2.3.0.44). Time-lapse imaging combined with z-stacks was performed every minute.

### Image analysis and Quantification

Image analysis and quantification were performed using Fiji ImageJ 1.52p (NIH) with the cell counter plugin for manual counting of GFP+ cells in zebrafish larvae from both sides.

### Statistical analysis

Statistical analysis was performed with GraphPad Prism 7 (Graphpad Software). A P value <0.05 was considered statistically significant. ****P<0.0001; ***P<0.001; **P<0.01; *P<0.05; ns (not significant) P ≥ 0.05.

## RESULTS

### Cytotoxicity of PD-L1.CAR.NK-92 cells against breast cancer cells *in vitro*

We first constructed a second-generation CAR containing the CD28 costimulatory and the CD3ζ activation domain with specificity for PD-L1 (PD-L1.CAR; **Figure 1A**), a checkpoint molecule frequently expressed in many solid tumors. The PD-L1.CAR was lentivirally transduced into the clinically applicable NK cell line NK-92 (PD-L1.CAR NK-92; **Figure 1B**), which has many similarities to primary NK cells and is a common model for CAR-NK cell studies (Klingemann, 2023; Maki et al., 2001). As a cancer model, we used the PD-L1-expressing metastatic breast cancer cell line MDA-MB-231 (**Figure 1C**). The PD-L1.CAR NK-92 cells showed strong CAR expression on the cell surface and exhibited efficient *in vitro* cytotoxicity against MDA-MB-231 cells (**Figure 1D,E**).

**Figure 1:**
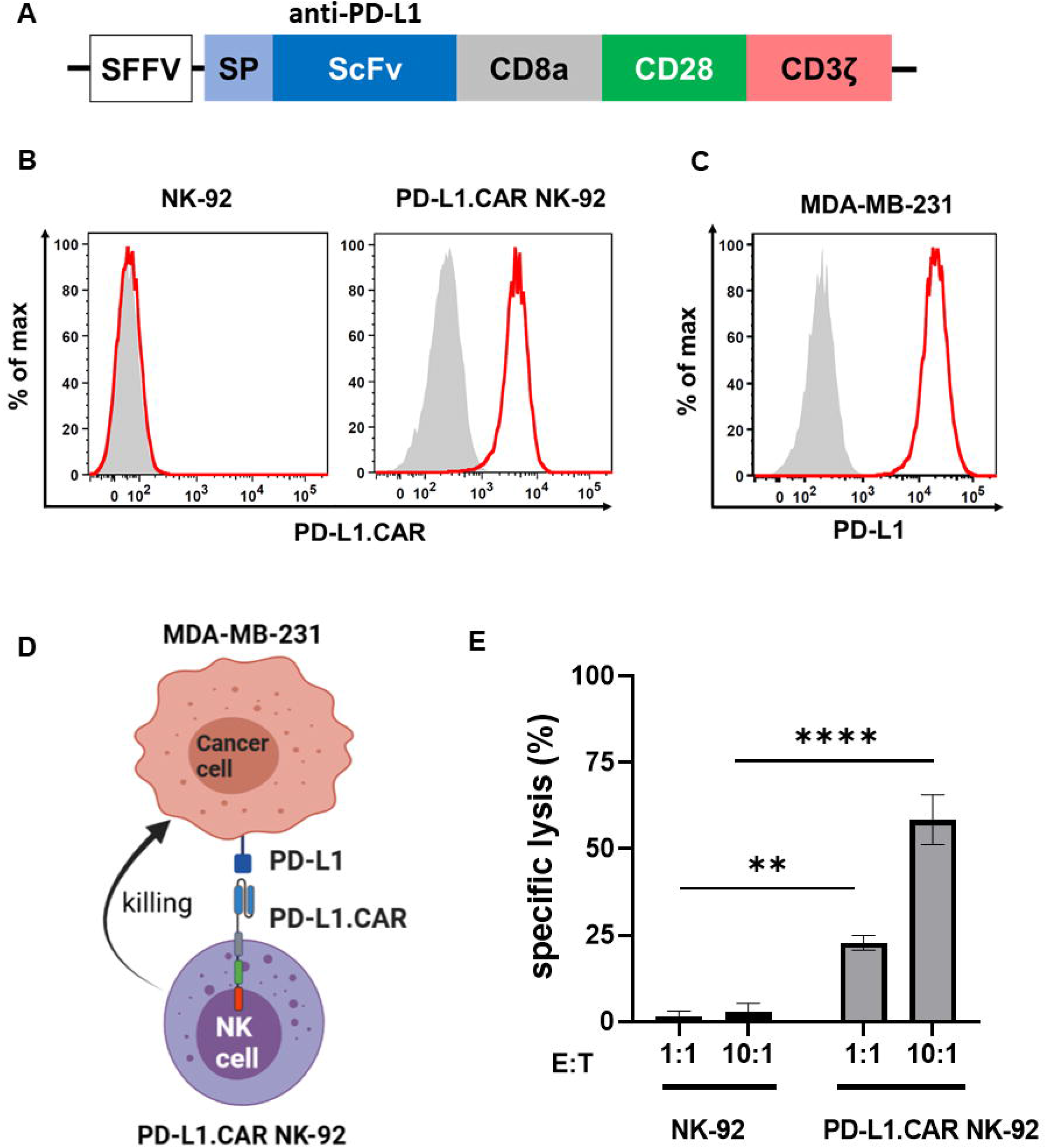
PD-L1.CAR NK-92 cells are highly cytotoxic against PD-L1+ targets *in vitro*. (A) Schematic representation of the PD-L1.CAR construct. A second-generation CAR under the spleen focus-forming virus (SFFV) promoter consists of PD-L1-specific scFv, CD8 hinge, CD28 transmembrane/costimulatory, and CD3ζ intracellular signaling domains. (B) Flow cytometric analysis of PD-L1.CAR expression on immunomagnetically enriched PD-L1.CAR NK-92 cells and parental NK-92 cells. PD-L1.CAR was detected using human recombinant PD-L1-Fc protein combined with anti-Fc secondary antibody. Filled gray areas indicate negative controls stained with secondary antibody only. (C) Flow cytometric analysis of PD-L1+ expression on the cell surface of MDA-MB- 231 cells. Representative data from at least 3 independent experiments are shown. (D) Illustration of the NK cell and cancer cell model. NK-92 cells were lentivirally transduced with PD-L1.CAR, and the PD-L1+ metastatic breast adenocarcinoma cell line MDA-MB-231 was used as a target. Created with BioRender. (E) PD-L1.CAR NK-92 or parental NK-92 cells were co-cultured with MDA-MB-231 cells at E:T ratios of 1:1 and 10:1 for 2 hours as indicated and specific *in vitro* cytotoxicity was measured by Europium-based cytotoxicity assay. One-way ANOVA with Tukey’s multiple comparisons was used to calculate statistics. Data were pooled from 3 independent experiments, and means ± SEM are shown.

### Injection of labeled cancer and CAR-NK cells into zebrafish embryos

To visualize the CAR-NK and cancer cells in the zebrafish xenograft model, we differentially labeled the effector and target cells. The MDA-MB-231 cells were stably transduced with GFP (MDA-MB- 231 GFP) without affecting PD-L1 expression (**Figure 2A**). The PD-L1.CAR NK-92 cells were labeled with the red membrane dye PKH26 prior to injection to distinguish them from the cancer cells (**Figure 2B**). We further confirmed that MDA-MB-231 GFP cells were efficiently killed by PD-L1.CAR NK-92 cells *in vitro*, with comparable efficacy compared to wild-type (WT) MDA-MB-231 target cells (**Figure 2C**). The MDA-MB-231 GFP cells were then injected into the circulation of zebrafish embryos via the DoC at 2 days post fertilization (dpf) using a micromanipulation device. Correctly injected embryos were sorted under the microscope, and 2.5 hours later red-labeled PD-L1.CAR NK-92 cells were injected into the DoC. Zebrafish embryos were then imaged 1 hour later. Both cancer and CAR-NK cells were clearly visible in the body of the zebrafish embryos. Cancer cells successfully migrated within the organism, especially to the tail region (including the CHT), head and ventral region of the zebrafish embryo (**Figure 2D**). Circulating CAR-NK cells were detected in the blood stream which is consistent with their subsequent injection, thus representing an optimal CAR-NK cell homing model to sites of diffuse metastatic cancer cell accumulation. With this approach, we established a novel assay for the visualization of CAR-NK cells and metastatic cancer cells in zebrafish embryos *in vivo*.

**Figure 2:**
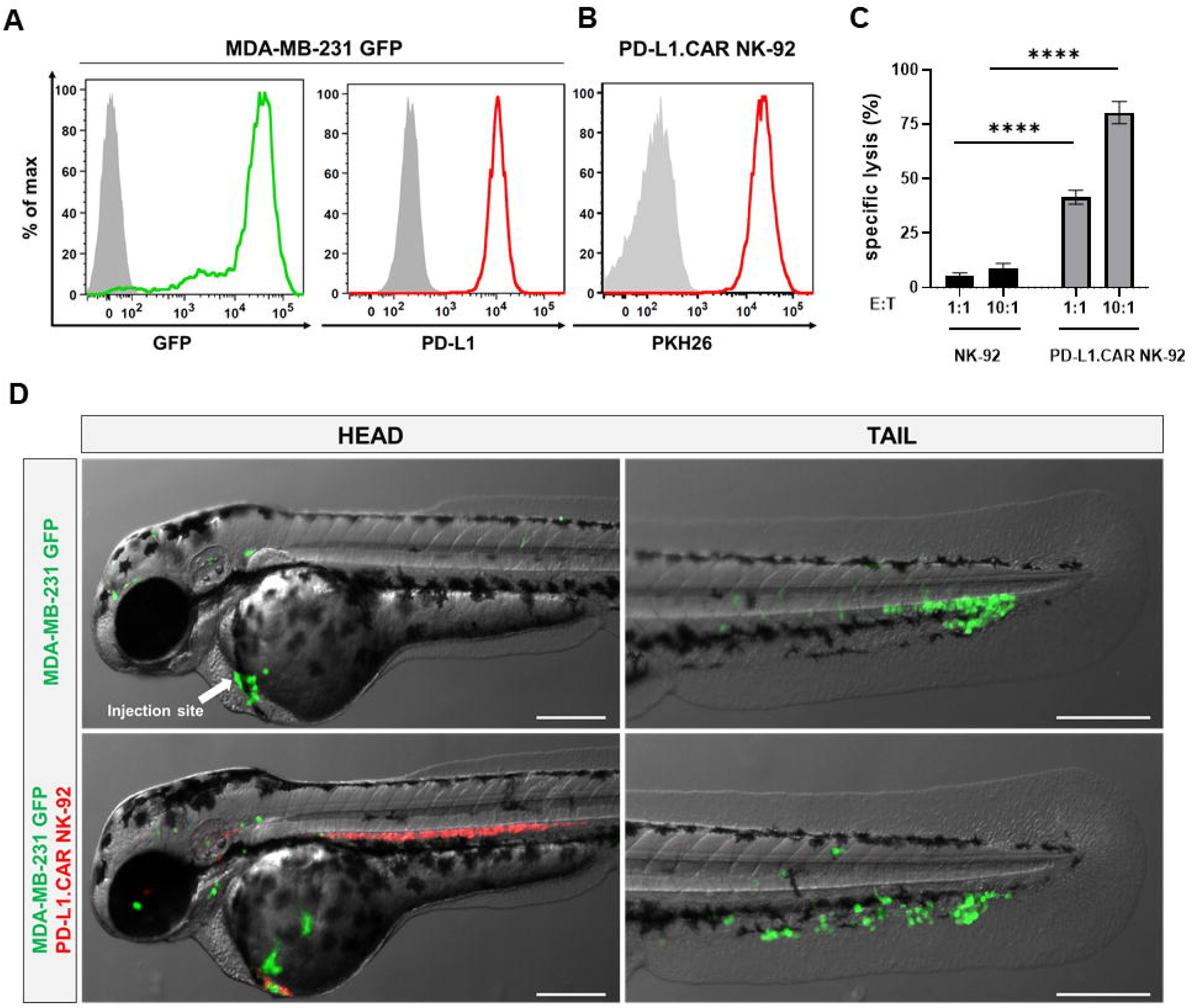
Labeled cancer and NK cells retain *in vitro* cytotoxicity and circulate in zebrafish larvae after injection. (A) MDA-MB-231 cells were transduced with GFP (MDA-MB-231 GFP). Expression of GFP and PD-L1 was analyzed by flow cytometry. Non-transduced and unstained cells shown in gray as negative controls for GFP and PD-L1, respectively. (B) PD-L1.CAR NK-92 cells were labeled with PKH26 and confirmed by flow cytometry. The filled gray histogram represents the unstained control. (C) PD-L1.CAR NK-92 or parental NK-92 cells were co-cultured with MDA-MB-231 GFP cells at E:T ratios of 1:1 and 10:1 for 2 hours as indicated, and specific *in vitro* cytotoxicity was measured by Europium-based cytotoxicity assay. Data were pooled from at least 3 independent experiments, and means ± SEM are shown. One-way ANOVA with Tukey’s multiple comparisons was used to calculate statistics. (D) MDA-MB-231 GFP cells were injected into the DoC alone (top) or injected at the same site 2.5 hours later with PD-L1.CAR NK-92 PKH26-labeled cells (bottom). Images of the head or tail region of zebrafish larvae were captured by fluorescence microscopy. Scale bar = 250 μm. Representative images are shown.

### Kinetics of *in vivo* cytotoxicity of PD-L1.CAR NK-92 cells in zebrafish

Next, we tested the *in vivo* cytotoxicity of the PD-L1.CAR NK-92 cells against the MDA-MB-231 GFP cells to identify the optimal time point for the final analysis. Therefore, we injected MDA-MB- 231 GFP cells into the DoC of a small cohort of zebrafish embryos followed by injection of red-labeled PD-L1.CAR NK-92 cells, unmodified parental NK-92 cells 2.5 hours later or without any follow-up cell injection (control). Individual zebrafish embryos were imaged immediately after NK cell injection and again at 24, 48 and 72 hours post-injection (hpi, **Figure 3A**). Cancer cells in the control group were distributed throughout the body of the zebrafish larvae, while in the NK-92 treated group, NK cells were found in close contact with the cancer cells. However, surviving cancer cells were detectable also at later time points (**Figure 3B**). In contrast, larvae treated with PD-L1.CAR NK- 92 cells were almost devoid of cancer cells at 48 and 72 hpi, while NK cells remained at the sites of interest. We quantified surviving cancer cells in the zebrafish across time points and confirmed significant *in vivo* cytotoxicity by PD-L1.CAR NK-92 cells compared to NK-92 and untreated controls as early as 24 hpi (**Figure 3C**). The cytotoxic effect was further enhanced at 48 and 72 hpi (10±5 and 5±3 remaining cells in PD-L1.CAR NK-92 compared to 39±6 and 36±5 remaining cells in NK-92, respectively). These data demonstrate that PD-L1.CAR NK-92 cells rapidly detect cancer cells *in vivo*, kill the majority of them within 24 hpi, and eliminated nearly all residual cancer cells within the following two days.

**Figure 3:**
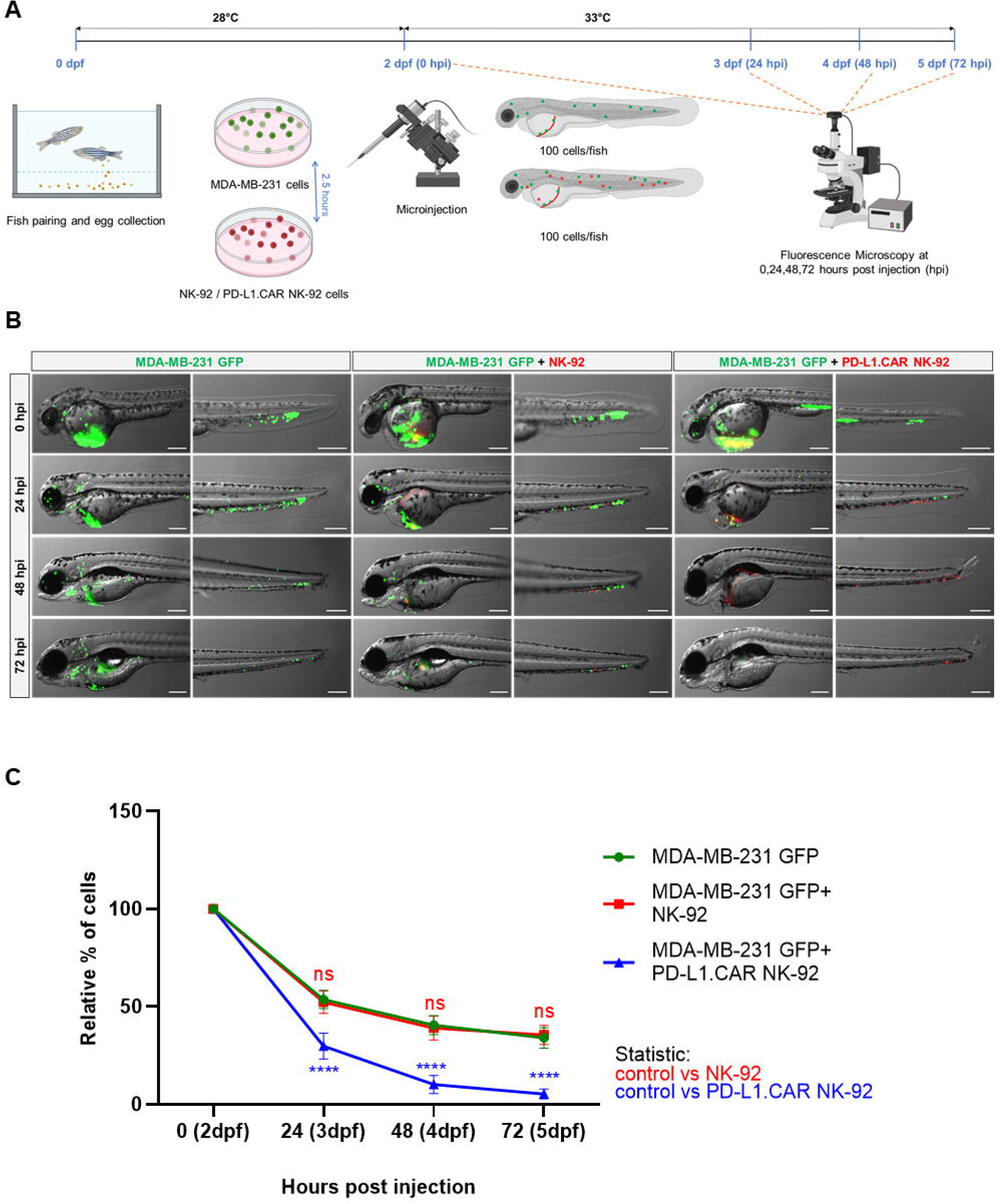
Kinetics of PD-L1.CAR NK-92 cytotoxicity in zebrafish *in vivo*. (A) Schematic time course of the experimental setup. Zebrafish larvae were injected with MDA-MB- 231 GFP cells at 2 days post fertilization (dpf) and 2.5 hours later with PKH26-labeled PD-L1.CAR NK-92 or parental NK-92 cells. For each cell type, 100 cells were injected on average. Images were captured by fluorescence microscopy at 24, 48, and 72 hours post injection (hpi). (B) Fluorescence micrographs showing zebrafish larvae injected with MDA-MB-231 GFP cells or combinations with NK-92 or PD-L1.CAR NK-92 at the indicated time points. MDA-MB-231 GFP cells are shown in green and PKH26-labeled NK cells are shown in red. Representative images are shown. Scale bar = 250 μm. (C) Quantification of MDA-MB-231 GFP cells throughout the zebrafish at the indicated time points and plotted as relative percentage of cells at 0 hpi. Data are pooled from 2 independent experiments and shown as mean ± SEM. n=3 (MDA-MB-231 GFP, MDA-MB-231 GFP+NK-92), n=4 (MDA-MB-231 GFP+PD-L1.CAR NK-92). Two-way ANOVA analysis with Tukey’s multiple comparison was used for statistical analysis.

### Evaluation of *in vivo* killing efficacy of PD-L1.CAR-NK and ErbB2.CAR-NK cells

To further validate the *in vivo* killing efficacy of PD-L1.CAR NK-92 cells in a pooled setup, we injected a larger cohort of zebrafish larvae with MDA-MB-231 GFP cells. Positively injected larvae were selected, divided into three groups and injected 2.5 hours later with PD-L1.CAR NK-92 cells, NK-92 control or left without any follow-up cell injection (control). Based on previous results, larvae were imaged at 48 hpi (**Figure 4A**). The quantification revealed that parental NK-92 cells exhibited low cytotoxicity with less than 50 % cancer cell reduction (23±16 remaining cells) compared to the control group (38±18 remaining cells). In contrast, PD-L1.CAR NK-92 cells almost completely eliminated cancer cells (7±7 remaining cells) in the majority of zebrafish larvae (**Figure 4B**). To test whether this system is applicable to other CAR-NK cell models, we used NK-92 cells with CAR targeting the ErbB2 (HER2) molecule (NK-92/5.28.z, here ErbB2.CAR NK-92) (Schonfeld et al., 2015). ErbB2 is overexpressed in many solid tumor cancer cells. To mimic this, we used the ErbB2- expressing breast cancer cell line MDA-MB-453 transduced with GFP (**Figure 4C**). Using this model we observed rapid killing of MDA-MB-453 GFP cells by ErbB2.CAR NK-92 cells as early as 8 hpi (18±22 remaining cells), whereas NK-92 cells mediated only moderate killing (47±28) compared to the control (96±36) (**Figure 4D**). Taken together, this demonstrates that CAR-NK cells with different lysis specificities can eliminate metastatic breast cancer cells in zebrafish larval xenografts.

**Figure 4:**
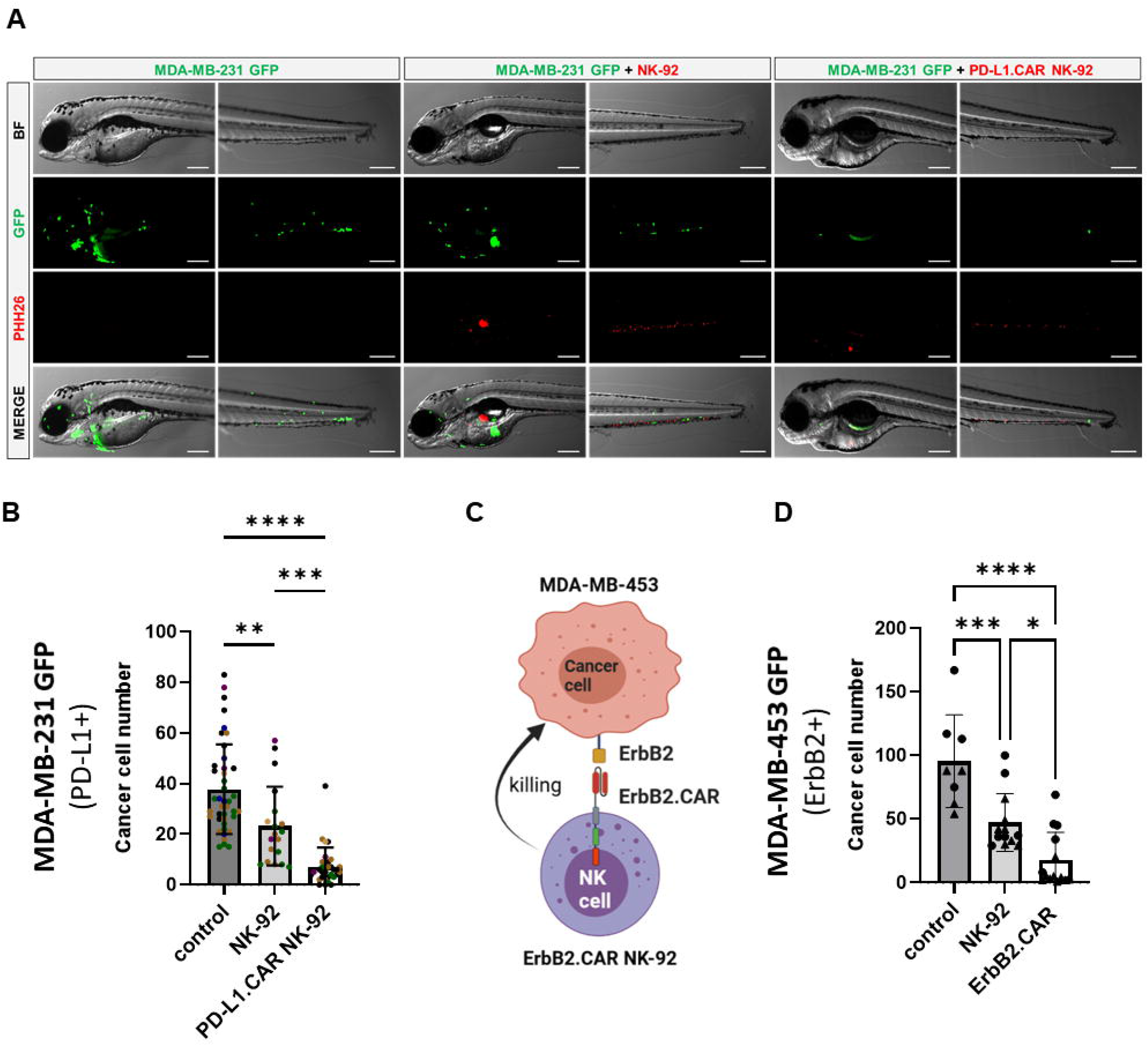
CAR NK-92 cells specific for PD-L1 or ErbB2 show high efficacy against resistant breast cancer cells in a zebrafish xenograft model. (A) Zebrafish larvae were injected at 2 dpf with MDA-MB-231 GFP cells and 2.5 hours later with PKH26-labeled PD-L1.CAR NK-92 or parental NK-92 cells. For each cell type an average 100 cells was injected. Images were captured by fluorescence microscopy at time points of 48 hpi. Scale bar = 250 μm. (B) The number of MDA-MB-231 GFP cells was quantified at 48 hpi. Data are pooled from 6 independent experiments and shown as mean ± SD. n=41 (MDA-MB-231 GFP), n=19 (MDA-MB-231 GFP+NK-92), n=31 (MDA-MB-231 GFP+PD-L1.CAR NK-92). (C) Schematic representation of ErbB2-specific CAR NK-92 (ErbB2.CAR NK-92) targeting ErbB2+ MDA-MB-453 metastatic breast cancer cell line. Illustration created with BioRender. (D) Zebrafish larvae were injected at 2 dpf with MDA-MB-453 GFP cells and 2.5 hours later with PKH26-labeled ErbB2.CAR NK-92 or parental NK- 92 cells. For each cell type, 100 cells were injected on average. Images were captured by fluorescence microscopy at 8 hpi and the number of cancer cells was quantified. Data were pooled from 2 independent experiments and means ± SD are shown. n=8 (MDA-MB-453 GFP) n=13 (MDA-MB- 453 GFP+NK-92), n=14 (MDA-MB-453 GFP+ErbB2.CAR NK-92). One-way ANOVA analysis with Tukey’s multiple comparison was used for statistical analysis. * p=0.0154.

### Live observation of CAR-NK cells killing tumor cells in the zebrafish xenograft

To further investigate the interaction of CAR-NK and cancer cells within the zebrafish host, we injected MDA-MB-231 GFP cells, followed by PD-L1.CAR NK-92 cell injection as described above, and performed confocal *in vivo* imaging of the zebrafish tail region 1 hour later (**Figure 5A**). The PD-L1.CAR NK-92 cells migrated to the cancer cells, conjugated with them, and subsequently killed the cancer cells (**Figure 5B; Supplementary video 1**). An important ability of CAR-NK cells is to migrate through the vasculature to reach metastatic cancer cells. To study this, we used the transgenic Tg(*flk1*:GFP) zebrafish strain with labeled endothelial cells (Beis et al., 2005). We observed that PD-L1.CAR NK-92 cells migrated to the disseminated metastatic cancer cells, demonstrating that CAR-NK cells can migrate through the zebrafish vasculature and identify individual metastatic cancer cells (**Figure 6, Supplementary video 2**). Taken together, our data demonstrate that zebrafish larvae are a suitable host to study the migration properties and efficacy of CAR-NK cells to eliminate metastatic cancer cells of solid tumor origin *in vivo*.

**Figure 5:**
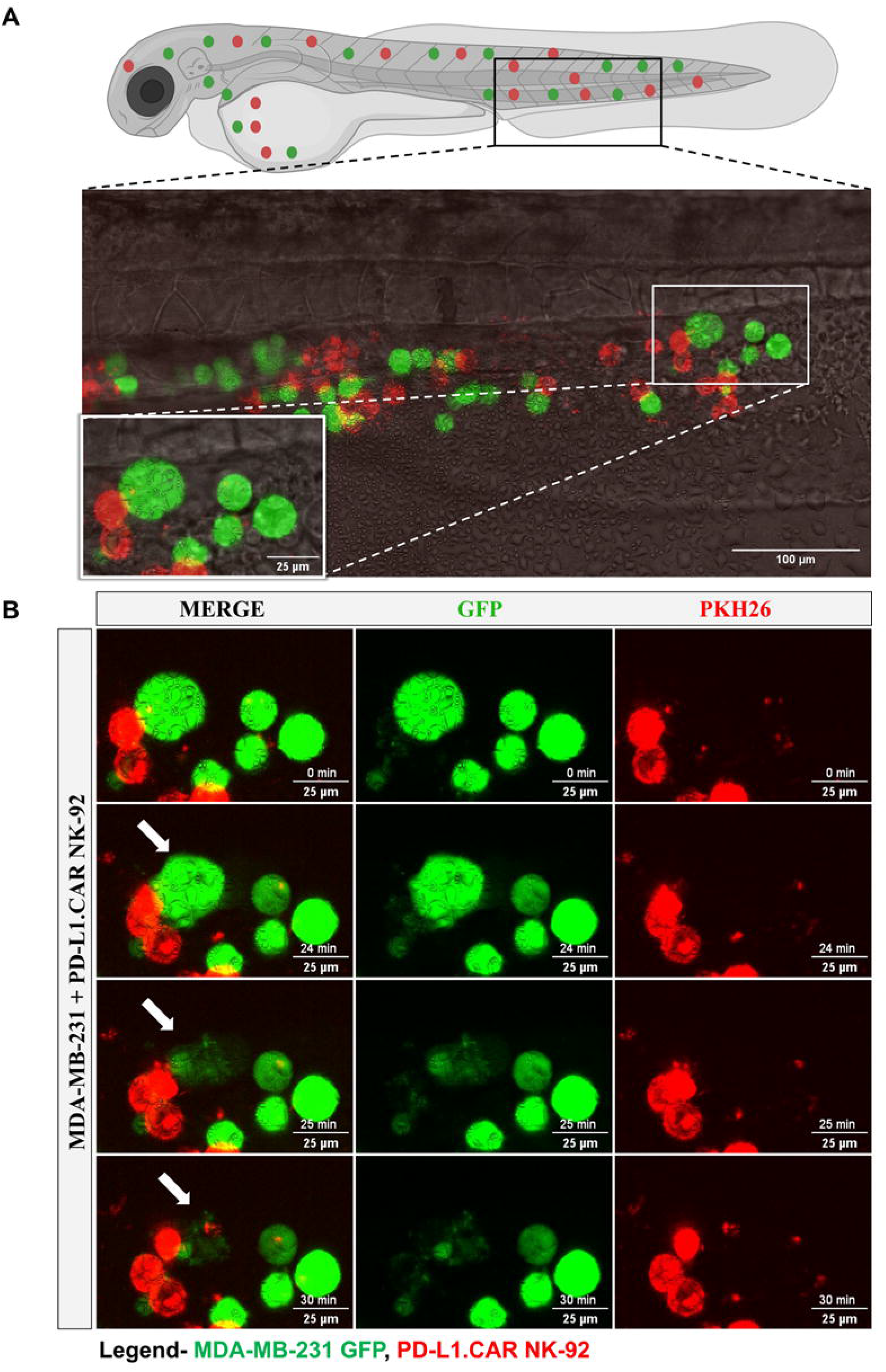
Time-lapse imaging of CAR-NK cell interaction with cancer cells and cytotoxicity in the zebrafish tail. MDA-MB-231 GFP cells and 2 hours later PD-L1.CAR NK-92 cells (PKH26-labeled) were injected into the DoC. One hour after injection, confocal microcopy images were taken in the tail region. (A) Representative picture showing the location of imaging. (B) Time-lapse images showing the interaction between NK cells (red) and cancer cells (green). The arrow indicates cancer cell killing. Scale bar = 25 μm.

**Figure 6:**
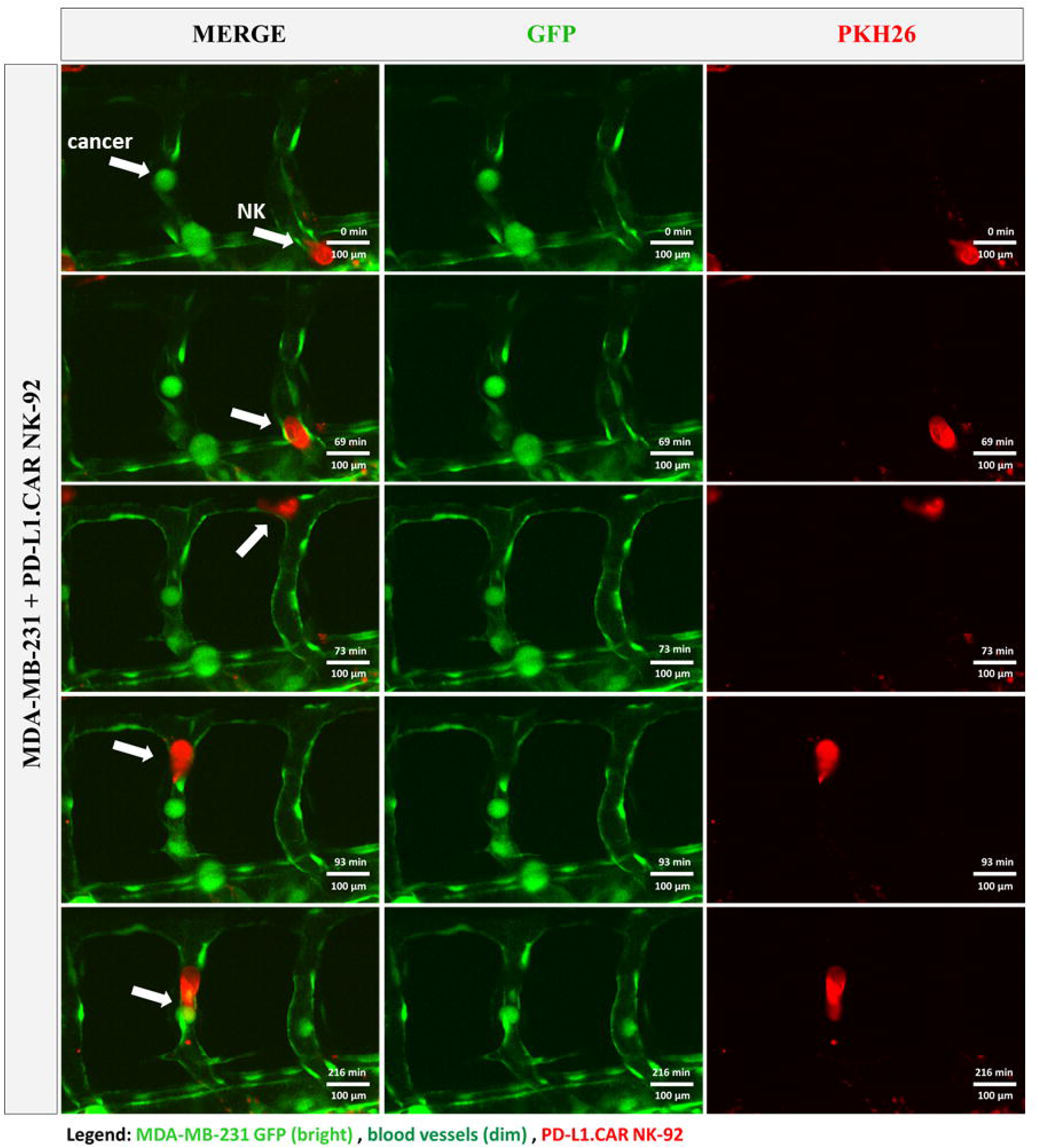
CAR-NK cells migrate to cancer cells within the vasculature. MDA-MB-231 GFP cells and PKH26-labeled PD-L1.CAR NK-92 cells were injected into transgenic Tg(*flk1*:GFP) zebrafish with endothelial cells expressing low levels of GFP. Confocal time-lapse microscopy shows NK cell migration to the metastatic breast cancer cell (indicated by arrows). Representative images are shown. Scale bar = 100 μm.

## DISCUSSION

CAR-NK cell therapies have recently emerged as a promising platform to treat otherwise incurable cancers without severe side effects. Despite promising clinical results in hematological malignancies, solid tumors remain challenging mainly due to their heterogeneity, resistance mechanisms, suppressive microenvironment, immune exhaustion and metastatic properties. Novel CAR or other targeted NK cell systems are rapidly emerging and therefore quick preclinical models to test their *in vivo* efficacy are required to facilitate the development of NK cell therapy. Currently, immunodeficient mouse xenografts are used, but they are limited in their use by high costs and long read out times.

Here, we describe a preclinical zebrafish xenograft model as an alternative and reliable system to test the efficacy of CAR-NK cells against metastatic breast cancer. Using two different CAR NK-92 systems targeting PD-L1 or ErbB2 molecules, we demonstrate that human NK cells survive in the zebrafish environment and retain specific cytotoxicity against metastatic breast cancer cells. Due to the absence of adaptive immune cells in zebrafish larvae, neither CAR-NK cells nor cancer cells appear to be rejected by the host´s endogenous immune system, although we cannot exclude that innate immune cells such as macrophages diminish CAR-NK and breast cancer cell numbers. In our metastatic model, we observed rapid killing of cancer cells in the periphery within 8 and 24 hours, respectively, depending on the target. The small size and transparency of the system allowed microscopic tracking of CAR-NK cells as well as of breast cancer cells in individual animals over time to obtain kinetic data on killing efficacy. The small size of the animals and high progeny number allowed us to inject larger cohorts of zebrafish, resulting in reliable results also in a pooled analysis setting. Our system will allow for medium-to high-throughput testing of novel CAR-NK cell products in early developmental stages of zebrafish before initiating time consuming mouse models, which in addition require the injection of a much higher cancer cell number. In this regard, the zebrafish system will be a promising tool to test the reaction of primary patient cancer cells to CAR-NK cell therapy. In fact, most cancer patients do not benefit from immunotherapy, and distinguishing responders from non-responders prior to treatment will allow for a reliable prediction of the treatment outcome. Because of the large number of available zebrafish embryos, low haltering costs and the small number of required patient cancer cells for injection (100-200 cells per embryo) the system is ideally suited for personalized clinical testing. Such a testing model allows fast recapitulation of the patient’s disease and testing of multiple CAR-NK cell products alone or in combination with other cell-based therapeutics such as CAR-T cells, whose action has recently been demonstrated in zebrafish embryos as a basis for T-cell based therapy (He et al., 2020; Pascoal et al., 2020). Of note, zebrafish xenografts also offer the possibility of testing the CAR-NK cells in combination with small drugs for synergistic effects, in analogy to previous studies using zebrafish for screening small drug libraries (Yan et al., 2019).

The zebrafish embryos’ small size and transparency allowed us to track CAR-NK cells and cancer cells at the single cell level, and to study CAR-NK/cancer cell interaction and cytotoxicity in real-time. In the future, confocal microscopy of cancer cell plus CAR-NK injected embryonic zebrafish could be used to study lytic granule polarization toward the immunological synapse, which is an important parameter associated with cancer cell resistance to NK cell cytotoxicity (Eitler et al., 2021). Furthermore, using transgenic zebrafish with a fluorescently labeled vasculature, we observe homing of metastatic cancer cells to the zebrafish periphery and intravascular migration of CAR-NK cells to these individual tumor cells, suggesting that CAR-NK cells can effectively find and eradicate circulating tumor cells and potentially micrometastases that are difficult to eliminate with standard therapies. In a clinical setting, these capacities can be exploited. For instance, the homing and persistence of novel improved CAR-NK cells including CIS (cytokine-inducible SH2-containing protein) knock out, transgenic IL-15 production and others can be tested in the zebrafish xenograft model (Delconte et al., 2016; Liu et al., 2018), as recently done to assess the efficacy of CAR-T cells expressing IL-15 and CCL19 (Zhou et al., 2022).

Although the used zebrafish xenograft model has many advantages, it is important to be aware of its limitations, one of which is its brevity itself. The model is very short-term and does not allow for the development of secondary (acquired) resistance mechanisms within the transplanted animal, as ultimately transplants might be rejected after development of innate immunity. However, recently an adult zebrafish model has been introduced that allows a minimum of 14 days of cancer-immune cell interaction albeit it is not clear whether this time is sufficient to study secondary resistance (Yan et al., 2021). Importantly, patient treatment needs to be started promptly in most cases, which is why short-term assays are generally desirable. Furthermore, the very first response to CAR-NK cell therapy is the first important parameter for healthcare professionals to apply the treatment. In line with this, we have shown that the CAR-NK cell efficacy can be measured as early as 24 hpi, but assessment can be extended to 72 hours if necessary. Similar to other xenograft animal models, the zebrafish do not contain human immune and stromal cells, limiting the ability to study complex immune cell interactions. This drawback can be overcome by co-injecting human stromal cells along with the cancer cells, as recently shown for CAR-T cells (He et al., 2020) or even by co-injecting immune cells from the patient. In order to enhance the survival of human cells in zebrafish, genetic engineering approaches introducing the production of human cytokines and growth factors can be taken, such as shown for human GM-CSF expression in zebrafish (Rajan et al., 2020). It is also important to note that although zebrafish have a wide temperature range due to temperature variation in their natural habitat, xenotransplanted individuals are usually maintained at a comparatively low temperature of 33 to 34°C, opposite to the standard 37°C of the mouse and human body. This could affect cancer and NK cell growth, proliferation and metabolism. However, we did not observe any effect of the lower temperature on CAR-NK cell killing efficacy, suggesting that this may be a minor issue. In addition, protocols for maintaining adult zebrafish at 37°C were recently established, suggesting that zebrafish xenografts can be further adopted for preclinical cell therapy models in the future (Yan et al., 2019).

In this work, we successfully assessed the efficacy of the CAR-engineered NK-92 cell line to eliminate breast cancer cells within xenotransplanted zebrafish larvae. NK-92 cells have many similarities with primary NK cells. Several other NK sources are currently being investigated as CAR effectors (Liu et al., 2020; Zhu et al., 2018) and further studies are required to test whether zebrafish xenografts can also be used for these cell types. In addition, NK cells have recently been shown to be successfully retargeted by various engagers such as monoclonal antibodies or bispecific antibodies, and the zebrafish model provides an attractive system for high-throughput *in vivo* testing of these drugs (Demaria et al., 2022; Pinto et al., 2022; Snyder et al., 2018; Vallera et al., 2021; Wingert et al., 2021). Therefore, we propose that the combination of novel engineered CAR-NK cells along with the versatility of zebrafish xenografts may facilitate the future advancement of CAR NK-based therapeutic approaches to the benefit of patients suffering from progressive disease.

## CONCLUSION

Here, we demonstrate that zebrafish embryos and larvae are a suitable *in vivo* system for testing the killing efficacy of CAR-NK cells against metastatic solid tumor cells. This system is flexible and will allow mid-throughput screening of new drugs at low cost, leading to faster development of highly effective CAR-NK cell products. It can also be refined to personalize testing of CAR-NK cell efficacy against patient-derived cancer cells and as a predictive tool to select the best-possible therapy for the patient.

### Supplementary Video 1

Video showing *in vivo* interaction and killing of MDA-MB-231 GFP cells by PD-L1.CAR NK-92 cells in the zebrafish larval xenograft model. Refers to Figure 5.

### Supplementary Video 2

Video showing migration of PD-L1.CAR NK-92 cells (red) in the transgenic zebrafish larval xenograft model with dim GFP vessels. The NK cell migrates within the circulation to the metastatic breast cancer cell (bright green). Refers to Figure 6.

## DECLARATIONS

## Acknowledgements

The authors would like to thank Erik Zenker and Madeleine Teichert for outstanding technical assistance, Hella Hartmann and Michael Gerlach for excellent assistance with the light microscopy facilities, Akash Emmanuel Aravindan for help with confocal data analysis. We are grateful to Katrin Lambert for teaching the microinjection procedure, to Alejandra Cristina López-Delgado for general support and to Marika Fischer, Daniela Mögel and Silvio Kunadt for excellent fish care. This work was supported by the Light Microscopy Facility of the TU Dresden CMCB Technology Platform. BioRender software was used for the graphical illustrations in the figures.

## Contributors

Designing of research studies: JE, FK, TT. Conducting experiments: NMS, POM, WR, AK. Acquiring data: NMS, AK. Analyzing data: NMS, JE. Writing the manuscript: JE, FK, TT, WSW, SRK. All authors read and approved the final version of the manuscript.

## Funding

This research was supported in part by the German Federal Ministry of Education (Clusters4Future SaxoCell, 03ZU1111DA) and German Red Cross Blood Donation Service internal grants to T.T., the German Research Foundation (DFG) through EI1223/2-1 to J.E., the DFG TRR67 (project 387653785), SPP2084 µBone (project KN 1102/2–1) and Transcampus (project tC2020_02_MED) to F.K. The work at the TU Dresden is co-financed with tax revenues based on the budget agreed by the Saxonian Landtag.

## Competing interests

TT and WSW are named as inventors on patents in the field of cancer immunotherapy owned by their respective institutions.

Other authors declare that they have no competing interests.

## Data availability statement

The original contributions presented in the study are included in the article/Supplementary Material. Further inquiries can be directed to the corresponding authors.

